# PACeR: a bioinformatic pipeline for the analysis of chimeric RNA-seq data

**DOI:** 10.1101/2022.05.25.493487

**Authors:** William T Mills, Andrew E. Jaffe, Mollie K Meffert

## Abstract

MicroRNAs (miRNAs) are small non-coding RNAs that function in post-transcriptional gene regulation through imperfect base pairing with mRNA targets which results in inhibition of translation and often destabilization of bound transcripts. Sequence-based algorithms historically used to predict miRNA targets face inherent challenges in reliably reflecting *in vivo* interactions. Recent strategies have directly profiled miRNA-target interactions by cross-linking and ligation of miRNAs to their targets within the RNA-induced silencing complex (RISC), followed by high throughput sequencing of the chimeric RNAs. Despite the strength of these direct chimeric miRNA:target profiling approaches, standardized pipelines for analyzing the resulting chimeric RNA sequencing data are not readily available. Here we present PACeR, a robust bioinformatic **p**ipeline for the **a**nalysis of **c**himeric **R**NA sequencing data. PACeR consists of two parts, each of which are optimized for the distinctive characteristics of chimeric RNA sequencing reads: first, read processing and alignment and second, peak calling and motif analysis. We apply PACeR to chimeric RNA sequencing data generated in our lab as well as a published benchmark dataset. PACeR has minimal computational power requirements and contains extensive annotation to broaden accessibility for processing chimeric RNA sequencing data and enable insights to be gained about the targets of small non-coding RNAs in regulating diverse biological systems.

## INTRODUCTION

MicroRNAs (miRNAs), discovered in 1993 (Lee et al. 1993; Wightman et al. 1993), are small non-coding RNAs (sncRNAs) that usually begin as primary miRNA transcripts (pri-miRNAs) transcribed by RNA polymerase II (Lee et al. 2002, 2004). Pri-miRNAs are processed in the nucleus by the microprocessor complex, comprised of Drosha and the double-stranded RNA binding protein DGCR8, to form a 60-70-nucleotide-long, stem-looped precursor miRNA (pre-miRNA) (Gregory et al. 2004; Han et al. 2004; Lee et al. 2003). Pre-miRNAs are exported from the nucleus after which the endoribonuclease Dicer cleaves the pre-miRNA stem to form a 22-base-pair RNA duplex (Ketting et al. 2001; Knight and Bass 2001). One strand of the RNA duplex is loaded onto an Argonaute protein (Ago), forming the RNA-induced silencing complex (RISC) (Miyoshi et al. 2005; Matranga et al. 2005), and directs the complex to target messenger RNAs by Watson-Crick-base-pairing with imperfect complementarity, usually within the mRNA 3’ untranslated region (3’ UTR). The RISC acts to inhibit protein synthesis by translation inhibition followed in most cases by transcript degradation (Fabian et al. 2009; Zdanowicz et al. 2009; Bazzini et al. 2012; Djuranovic et al. 2012; Béthune et al. 2012; Mishima et al. 2012; Moretti et al. 2012; Meijer et al. 2013; Ricci et al. 2013), though the order of these events is under some debate (Hu and Coller 2012). As most mammalian mRNAs are targets of miRNAs (Friedman et al. 2009), the potential for miRNAs to coordinately influence gene expression and impact cell biology is immense. In addition to miRNAs, other sncRNAs such as tRNA fragments (tRFs) have also been shown to associate with Ago and mediate post-transcriptional gene regulation (Kumar et al. 2014; Kuscu et al. 2018; Guan et al. 2020; Xiao et al. 2021; Zuo et al. 2021; Wang et al. 2022).

Several bioinformatic tools have been developed to predict the mRNA targets of miRNAs including TargetScan (Lewis et al. 2003) (and later TargetScanS (Lewis et al. 2005)), DIANA-microT (Kiriakidou et al. 2004), miRanda (John et al. 2004), PicTar (Krek et al. 2005), and miRwalk (Dweep et al. 2011), as well as others for specific organisms such as *Drosophila* (Enright et al. 2003; Stark et al. 2003). While a variety of parameters have been used to make target predictions, the algorithms mainly rely on combining principles such as evolutionary conservation of the miRNA and its binding site within mRNA 3’ UTRs, binding free energy of the miRNA target duplex, degree of complementarity between nucleotides 2-7 from the 5’ end of the miRNA (the miRNA “seed” region (Lewis et al. 2003)) and its target, size and position of mismatches/bulges across the miRNA-target duplex, and combinatorial effects of multiple miRNA binding sites within close proximity. While these methods refine a list of potential miRNA target sites, many features of cell and miRNA biology can cause these predictions to be inadequate and to contain false positives and false negatives: (1) noncanonical seed interactions such as those with G-U wobbles, bulging, and mismatched bases (Lal et al. 2009; Helwak et al. 2013); (2) base-pairing outside of the miRNA seed influencing target specificity (Broughton et al. 2016; Chipman and Pasquinelli 2019; Duan et al. 2022); (3) miRNA binding sites outside 3’ UTRs such as in coding regions (Tay et al. 2008; Forman et al. 2008) and 5’ UTRs (Lytle et al. 2007); (4) RNA binding proteins and mRNA secondary structure impacting miRNA binding site accessibility (Ciafrè and Galardi 2013; Long et al. 2007); (5) post-translational modifications of Argonaute and various co-factors altering miRNA function (Hammell et al. 2009; Johnston and Hutvagner 2011); and (6) subcellular localization of miRNAs and potential targets (Wang and Bao 2017). Further, while altered miRNA abundance is a common feature of both disease and physiological responses, bioinformatic algorithms do not accurately predict the consequences of altered miRNA levels on target repression. Taken together, these factors highlight the need for broad adoption of less ambiguous genome-wide methods for determining miRNA-target interactions by RNA scientists.

The first attempt to resolve the ambiguity of determining miRNA-target interactions utilized crosslinking and immunoprecipitation (CLIP) to identify miRNAs and mRNAs bound directly to Ago using high-throughput sequencing. This method, called “Ago HITS-CLIP,” improved the accuracy of predicted miRNA-target interactions by only considering RNAs bound directly to Ago (Chi et al. 2009). However, this method still relied on bioinformatics to predict which Ago-bound miRNAs and mRNAs might be interacting. To address this ambiguity, several groups have developed methods incorporating an intermolecular ligation step to directly ligate the miRNA to its target within the RISC (Table 1) (Helwak et al. 2013; Grosswendt et al. 2014; Moore et al. 2015; Broughton et al. 2016). Since the miRNA and its *in vivo* cross-linked mRNA target are ligated together into a single chimeric RNA molecule, targets no longer need to be predicted by algorithms because this information can be ascertained directly from downstream high-throughput sequencing reads. Multiple strategies, as outlined in Table 1, have now utilized variations of this chimeric RNA approach followed by sequencing for investigating miRNA-target interactions across biological models and experimental settings.

**Table 1.** Biochemical methods for profiling miRNA-target interactions.

| Method | Reference | Model System | Photoactivatable Nucleotides | Argonaute | Chimeric RNAs |
| --- | --- | --- | --- | --- | --- |
| Argonaute HITS-CLIP (Ago HITS-CLIP) | (Chi et al. 2009) | Mouse (P13 neocortex) | No | Endogenous | No |
| Photoactivatable-Ribonucleoside-Enhanced Crosslinking and Immunoprecipitation (PAR-CLIP) | (Hafner et al. 2010, 2012) | Human (HEK293 cells) | 4-thiouridine | FLAG/HA-tagged | No |
| Modified Individual-nucleotide resolution UV-Cross-Linking and Immunoprecipitation (iCLIP) | (Broughton and Pasquinelli 2013) | *C. elegans* | No | Endogenous | No |
| Cross-Linking, Ligation, and Sequencing of Hybrids (CLASH) | (Helwak et al. 2013) | Human (HEK293 cells) | No | Protein A-TEV cleavage site-His6 tripartite-tagged | Yes |
| Modified *in vivo* PAR-CLIP (iPAR-CLIP) | (Grosswendt et al. 2014) | *C. elegans* | 4-thiouridine | Endogenous | Yes |
| Covalent Ligation of Endogenous Argonaute-bound RNAs-CLIP (CLEAR-CLIP) | (Moore et al. 2015) | Mouse (P13 neocortex)<br>Human (Huh-7.5 cells) | No | Endogenous | Yes |
| Argonaute iCLIP (AGO iCLIP) | (Broughton et al. 2016) | *C. elegans* | No | Endogenous | Yes |
| Quick CLASH (qCLASH) | (Gay et al. 2018) | Human (TIVE-LTC cells) | No | Endogenous | Yes |

The scientific value of removing the ambiguity of miRNA-target prediction has already produced significant insights about miRNA function and has the potential to shed light on the functions of other sncRNAs as well (Helwak et al. 2013; Grosswendt et al. 2014; Moore et al. 2015; Broughton et al. 2016; Gay et al. 2018; Kumar et al. 2014). However, published information about how the sequencing reads from chimeric RNAs are processed bioinformatically is typically incompletely described, lacks uniformity, and uses tools for data processing that are often cumbersome to apply or are out of date. These data processing difficulties are a barrier to the more widespread adoption of chimeric RNA approaches for understanding miRNA-target interactions, as well as potentially impacting the rigor and reproducibility of published findings. Here we introduce PACeR, a well-annotated **p**ipeline for the **a**nalysis of **c**himeric **R**NA sequencing data, that allows investigators to readily process high-throughput sequencing data from chimeric RNAs derived from a variety of molecular approaches and biological systems, including those described above (Table 1). Since there is no one set of parameters that will satisfy the needs of all experiments, providing detailed annotation with demarcated decision points and utilizing well-supported bioinformatics tools, rather than relying on scripts and packages developed by individual groups, is critical for allowing this pipeline to be readily adapted for users’ specific needs over sustained usage. In addition, the pipeline is annotated to aid as a tutorial for users with modest levels of bioinformatics experience. To further facilitate broad adoption, this pipeline has been developed to function efficiently on most personal computers, removing any necessity for high performance computer clusters or cloud computing.

## RESULTS AND DISCUSSION

### Limitations of current pipelines for analysis of chimeric miRNA:target RNA-seq data

Published biochemical approaches to produce chimeric RNAs (e.g. Table 1) each provide some description of how RNA sequencing reads were processed. However, providing streamlined protocols is often not the focus and stand-alone details required to reproduce RNA processing procedures can be scarce. Some publications indicate that data are processed “as described” in prior work and include only the additional parameters used in the current publication without including the analysis code. In addition, most publications describing the processing of chimeric RNA reads have used analysis packages that are not regularly updated or were replaced post-publication, such as the Crosslinking Induced Mutation Site (CIMS) software package (Zhang and Darnell 2011; Moore et al. 2014) that was replaced with the CLIP Tool Kit (CTK) (Shah et al. 2017). A further barrier to adoption by other laboratories is that some previously utilized analysis packages are limited in their ability to process chimeric reads generated from the diverse sequencing library preparation methods. For example, a script may specify removal of 5’ sequencing barcodes (e.g. CTK package) without an option for removing the 3’ barcodes in chimeric reads which are generated in other preparations. In other instances, features of bioinformatic pipelines, such as accession by online servers (including Galaxy) or wrapping of command line functions into other programming languages (such as R), may preclude the ability of users to manipulate key parameters for specific analysis needs. This limitation can include, for example, not allowing user specification of a reference genome aligner for read-mapping that suits their purposes (e.g. a splice-aware aligner or an aligner requiring less RAM). The use of wrapper functions to move command line functions into other programming languages can also have the unintended consequence of making the code cumbersome, requiring more arguments to accomplish the same task, and relying on the documentation of the wrapper function rather than documentation of the analysis function. Further, scripts coded in bash (a UNIX shell and command language) that perform modular steps have greater flexibility and readability for the typical user compared to end-to-end processing scripts written in programming languages such as perl. The bioinformatic pipeline presented here makes use of easy-to-use bash scripts that can be readily adapted for users’ specific needs by following the provided annotation and adjustable parameters.

### Pipeline overview

We developed a bioinformatic pipeline for analysis of chimeric RNA sequencing data (PACeR) which emphasizes simplicity and stability, annotation, and modifiability (Figure 1). The pipeline consists of two general parts: <u>first</u>, read processing and alignment and <u>second</u>, peak calling and motif analysis. Command-line scripts perform each of the requisite steps in modular fashion using intentionally straightforward code to allow accessibility for users with minimal bioinformatics experience. Transparency regarding how data are being processed is provided by thorough annotation throughout the scripts explaining the use of parameters and references to the user manuals for specific packages. Adjustable parameters are highlighted to allow for the pipeline to be readily adapted for users’ needs. The external packages used in this pipeline (e.g. FastQC, MultiQC, FLASh, Cutadapt, BLAST, HISAT2, Samtools, GATK, and BEDTools) were selected based on factors such as ease of use, thoroughness of documentation, frequency of use in publications, and degree of support (i.e. update frequency). This approach is intended to enhance the reproducibility of results across users and to facilitate user customization of the analysis for their unique purposes. Below, we proceed through a tutorial of the steps and considerations within the PACeR analysis pipeline while highlighting some areas of potential parameter customization.

**FIGURE 1.**
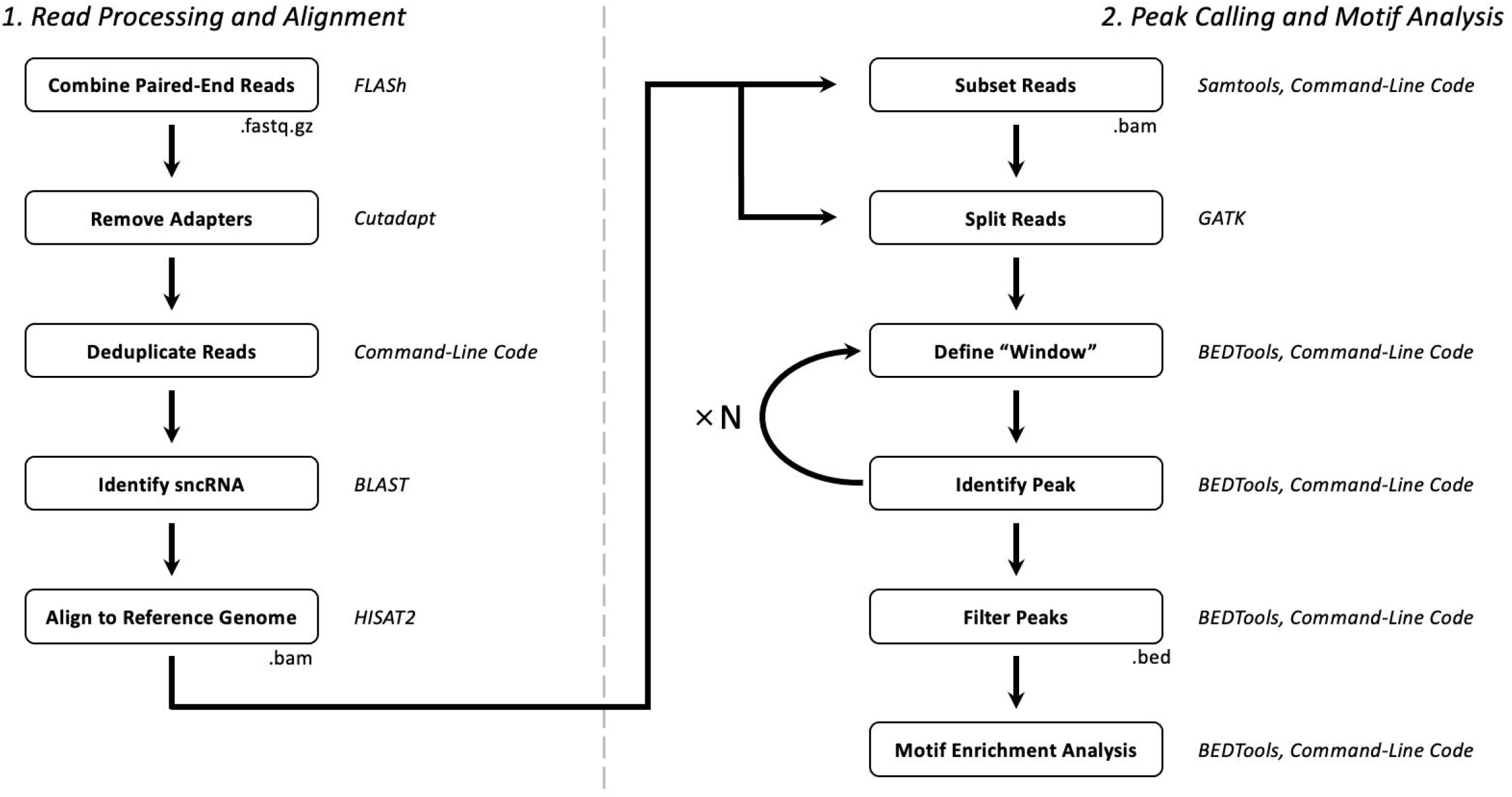
Overview of PACeR workflow. (1) Read Processing and Alignment: Compressed FASTQ files are combined using FLASh followed by adapter removal using Cutadapt, deduplication using command-line code, scnRNA identification using BLAST, and alignment to the reference genome using HISAT2 to produce BAM files. (2) Peak Calling and Motif Analysis: BAM files are subset by individual sncRNAs or sncRNA families or processed as a whole before splitting reads across unmapped regions using GATK, identifying “windows” of read coverage and peaks within “windows” using BEDTools and command-line code (for a user-defined number of cycles), and filtering peaks by coverage across libraries using BEDTools and command-line code. If reads were subset by sncRNAs or sncRNA family, motif enrichment analysis can be performed to determine the most prevalent 6-mer or 7-mer motifs within peaks. Italicized text indicates packages or toolkits used in the corresponding step.

#### Quality Control

The quality of the sequencing data is first assessed using FastQC (Andrews 2010) and visualized with MultiQC (Ewels et al. 2016). FastQC is one of the most broadly used bioinformatic tools for assessing the quality of high-throughput sequencing data and includes parameters such as per base and per sequence quality scores, sequence length distribution, sequence duplication levels, and adapter content. As nearly all sequencing experiments incorporate multiple samples, it is useful to visualize the quality of each sample side-by-side. MultiQC allows for multiple FastQC reports to be combined into a single HTML file and visualized together so that variations between samples can be discerned more readily.

##### The unique nature of the chimeric RNAs being sequenced can cause data to be flagged for quality concerns

during processing. For example, the length of the chimeric RNAs is quite variable, particularly at the 3’ end of the target RNA, so these inserts may be shorter than the read length of the sequencing reaction. This results in forward and reverse reads that contain the 3’ and 5’ adapter sequences, respectively. An overabundance of adapter sequence in reads is flagged by FastQC as low-quality data but in reality, this is simply an artifact of the inherent properties of these chimeric RNAs. Table 2 summarizes several of the FastQC parameters that may be negatively impacted by the intrinsic properties of sequenced chimeric RNAs but that are not necessarily indicative of low-quality data. While these factors explain some of the potential reasons for low quality scores, they do not exclude the possibility of real issues with the data that should be considered before proceeding.

**Table 2.** FastQC quality parameters that may be impacted by the nature of chimeric RNAs.

| FastQC Parameters | Explanation of low score |
| --- | --- |
| Per base sequence content | The patterned nature of the chimeric RNAs (e.g miRNA followed by mRNA) can cause a non-random collection of bases to occur throughout the read such as the overabundance of uracil at the first base of the miRNA and adenosine at the base immediately downstream of the seed binding site in the target RNA. |
| Per sequence GC content | The patterned nature of the chimeric RNAs (e.g miRNA followed by mRNA) can cause a non-random collection of bases to occur throughout the read such as the overabundance of uracil at the first base of the miRNA and adenosine at the base immediately downstream of the seed binding site in the target RNA. |
| Sequence duplication | The complexity of the library preparation protocols can result in excess primers and adapters being present in the sequencing reaction. |
| Overrepresented sequences | The abundance of chimeric RNAs that are shorter than the read length of the sequencing reaction will result in a higher-than-expected abundance of adapter sequence within reads.<br>The complexity of the library preparation protocols can result in excess primers and adapters being present in the sequencing reaction. |
| Adapter Content | The abundance of chimeric RNAs that are shorter than the read length of the sequencing reaction will result in a higher-than-expected abundance of adapter sequence within reads. |

#### Combining Paired-End Reads

Paired-end reads are combined using FLASh (Magoc and Salzberg 2011). This helps ensure that both the 5’ and 3’ adapters are present which improves confidence that the resulting read represents successful processing of the chimeric RNA molecule with the random nucleotide barcodes at either end of the chimeric RNA. Note that chimeric RNAs that are shorter than the read length (e.g. for a standard selected sequencing read length of 150 bp) can cause paired-end reads to exist in what is termed the “outie” orientation rather than the canonical “innie” orientation (described at http://gensoft.pasteur.fr/docs/FLASH/1.2.11/flash). As a result, when run with the default settings, FLASh may return the following error message:

> “WARNING: An unexpectedly high proportion of combined pairs (16.72%) overlapped by more than 65 bp, the --max-overlap (-M) parameter. Consider increasing this parameter. (As-is, FLASH is penalizing overlaps longer than 65 bp when considering them for possible combining!)”

This issue can be resolved using the `-O (--allow-outies)` and `-M 150 (--max-overlap=150)` parameters. The maximum overlap (150) in this case is selected based on the 150 bp read length of our representative data and could be adjusted.

#### Adapter Removal

Adapter sequences are removed using Cutadapt (Martin 2011). To improve confidence that a read represents successful processing of the chimeric RNA molecule and inclusion of the random nucleotide barcodes, only reads containing both the 5’ and 3’ (linked) adapters are considered (-g CTACAGTCCGACGATC…TGGAATTCTCGGGTGCCAAGG). For application to data without both 5’ and 3’ adapters (e.g. single-end sequencing), these parameters can be modified according to Cutadapt’s documentation (https://cutadapt.readthedocs.io/en/stable/guide.html) to not require presence of both adapters (e.g. using the `-a` and `-g` command-line options to denote the 3’ and 5’ adapters, respectively). Cutadapt is also used to trim low quality bases from the ends of reads using the `-q 30` (`--quality-cutoff 30`) option and to set a minimum length using the `-m 30` (`--minimum-length 30`) option.

#### Deduplication

While library preparation protocols should minimize PCR amplification cycles, PCR duplicates must be removed prior to downstream analyses to enable quantitative comparisons of sncRNA:target RNA interactions between samples. PCR duplicates and multiple occurrences of the same sncRNA:target interaction are distinguished using the random nucleotides included in the 5’ and/or 3’ adapters that are ligated to the chimeric RNA during library preparation in our protocol. Exact sequence duplicates are removed and the number of PCR duplicates for a given read is appended to the read ID so that duplication rates of chimeric sequences may be analyzed.

#### Barcode Removal

The random nucleotide barcodes at the 5’ and/or 3’ ends of chimeric RNA are removed and appended to the read ID. The length of these barcodes can be defined by the user to allow for flexibility with different biochemical protocols.

#### sncRNA Identification

The identity of the sncRNA within the chimeric RNA is determined with the widely-utilized BLAST program (Altschul et al. 1990) by aligning chimeric reads to a reference FASTA file. In this PACeR tutorial, the given reference FASTA file consists of miRNAs (from miRBase (Kozomara et al. 2019); https://www.mirbase.org/) and tRFs (from tRFdb (Kumar et al. 2015); http://genome.bioch.virginia.edu/trfdb/), however, the reference file can be compiled by users according to their needs. BLAST results can be filtered using user-defined parameters; here, BLAST results were filtered requiring a minimum alignment of 14 bases, an e-value less than 0.05, no more than 2 mismatches, no more than 1 gap, and no gaps if 2 mismatches are present.

#### sncRNA-first Chimeras

The biochemical steps for generating chimeric RNAs strongly biases the reaction in the 5’-sncRNA:mRNA-3’ orientation (sncRNA-first), although at a low frequency chimeras may be generated in the 5’-mRNA:sncRNA-3’ orientation (sncRNA-last) (Moore et al. 2015). Only chimeras in which the sncRNA starts at the first base within the read following adapter and barcode removal are considered for downstream analyses. Further, users can also define the minimum length of bases downstream of the sncRNA; here, the minimum length was set at 15 bases. The minimum length defined by the user should be selected with consideration for the requirements of the genome aligner used in the subsequent step. Once the sncRNA-first chimeras have been identified, the sncRNA sequence is removed from the read and the sncRNA name is appended to the read ID.

#### Target Identification

The sequence downstream of the sncRNA is aligned to the mouse genome using the splice-aware aligner HISAT2 (Kim et al. 2019). While other aligners may be used (e.g. Bowtie, STAR, Novogene, etc.), HISAT2 is reported to match the accuracy of other aligners (Baruzzo et al. 2017) and requires substantially less RAM than aligners like STAR. This allows users to run PACeR on personal computers without the need for access to computing clusters. To improve confidence in downstream analyses, only uniquely aligned reads (those that align to the genome once and only once) are considered.

#### Peak Calling

The calling of peaks, regions in which multiple reads from a particular sncRNA or sncRNA family overlap, is used to identify confidently targeted regions of the genome, to search for sncRNA binding motifs, and to aid in making comparisons across biological samples. Traditional peak callers such as PureCLIP (Krakau et al. 2017), Homer (Heinz et al. 2010), and MACS (Zhang et al. 2008) are usually designed for experiments that generate a normal distribution of sequencing reads around sites in the genome such as CHIP-seq or RNA binding protein CLIP-seq. These peak callers perform less well with data generated from the chimeric RNA strategies discussed here, because these approaches generate a non-normal distribution of reads. This skewed distribution of reads can occur due to the biochemical strategy of sncRNA:target RNA ligation since there are limits on the length of the 5’ end of the target RNA outside the Ago complex that will allow for successful ligation to the sncRNA and no such limits on the length of the 3’ end of the target RNA outside the Ago complex (Figure 2A). This produces a skewed distribution of reads around sncRNA binding sites in the genome that, when visualized, appears as peaks that shoulder toward the 3’ end of the gene (Figure 2B). Additionally, traditional peak callers are often unable to detect multiple binding sites within close proximity to one another, binding sites within introns, and binding sites within noncoding RNAs. Each of these scenarios, and the ability of PACeR to resolve them, are presented below in the results section for the evolutionarily conserved let-7 family of miRNAs.

**FIGURE 2.**
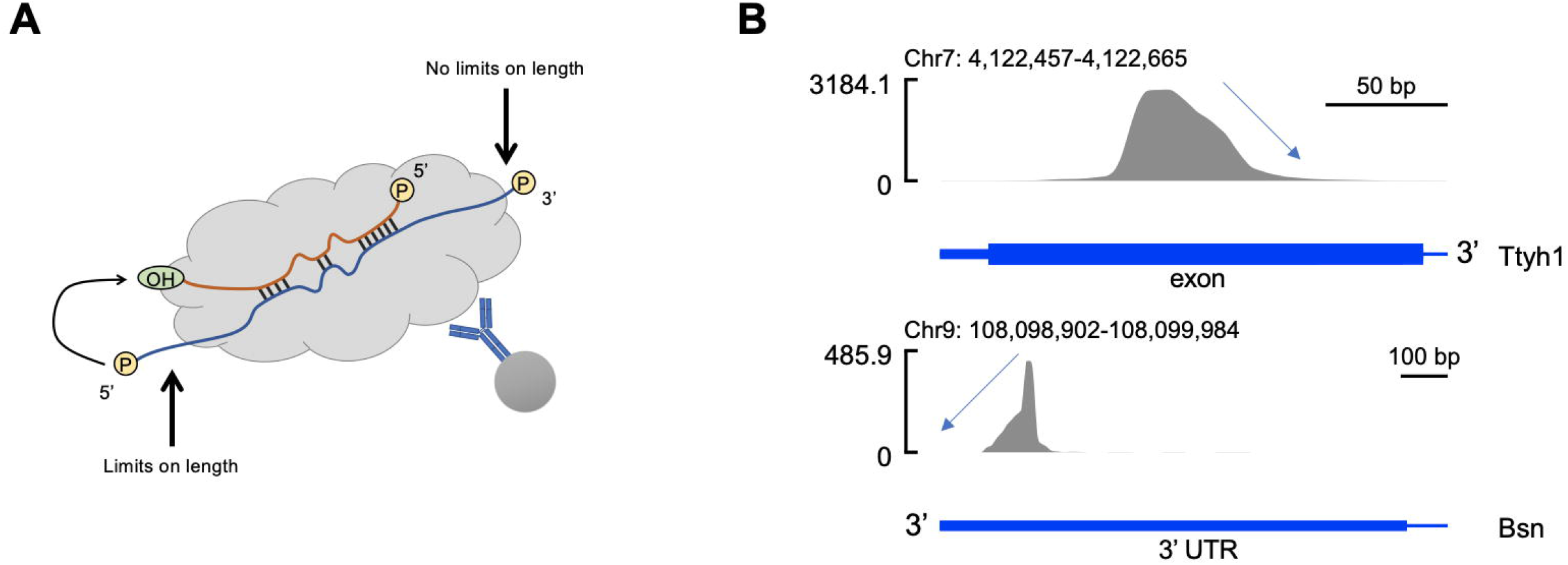
Differential length restrictions on the 5’ and 3’ ends of the target RNA during the generation of chimeric RNAs results in a non-normal distribution of reads around sncRNA binding sites. **(A)** Since the ligation reaction predominantly occurs in the 5’-sncRNA:target-RNA-3’ direction, the length at the 5’ end of the target RNA must be within an acceptable range of lengths in order for the ligation to be successful. Since the ligation is usually not occurring at the 3’ end of the target RNA, no such length restriction exists. **(B)** The length restrictions at the 5’ end of the target RNA and lack of restriction at the 3’ end results in peaks that shoulder toward the 3’ end of the gene. Representative data are chimeric reads containing let-7 family miRNAs. Blue bars are graphical representations of gene regions. Peaks visualized using SparK (Kurtenbach and Harbour 2019).

For peak calling, reads from all samples are split across unmapped regions using the Genome Analysis Toolkit (GATK) (Auwera and O’Connor 2020), converted to BED format using BEDTools (Quinlan and Hall 2010), and combined into a single file. The combined BED file is modified by appending the sncRNA name to the chromosome number; this enhances the computational efficiency of PACeR by allowing peaks for all sncRNAs to be called at once without having to create separate BED files for every sncRNA. Overlapping reads are merged using BEDTools and regions of the genome with “significant” read coverage, termed ‘windows’, are identified. In our analyses we only consider regions with three or more reads of coverage as “significant,” however, this parameter can be changed by the user. The region of maximum coverage within the window is then determined and termed the ‘primary peak’. Due to the potential for multiple sncRNA binding sites to be in close proximity (i.e. within the same window), reads that fall under the primary peak are removed and the process is repeated by identifying new windows and then finding the region of maximal coverage within the new window in an iterative fashion (Figure 3); the number of times this process is repeated can be adjusted by the user as dictated by their data. Once the peaks have been identified, they are filtered to require that each peak be supported by reads from at least 2 libraries; this parameter can also be adjusted by the user.

**FIGURE 3.**
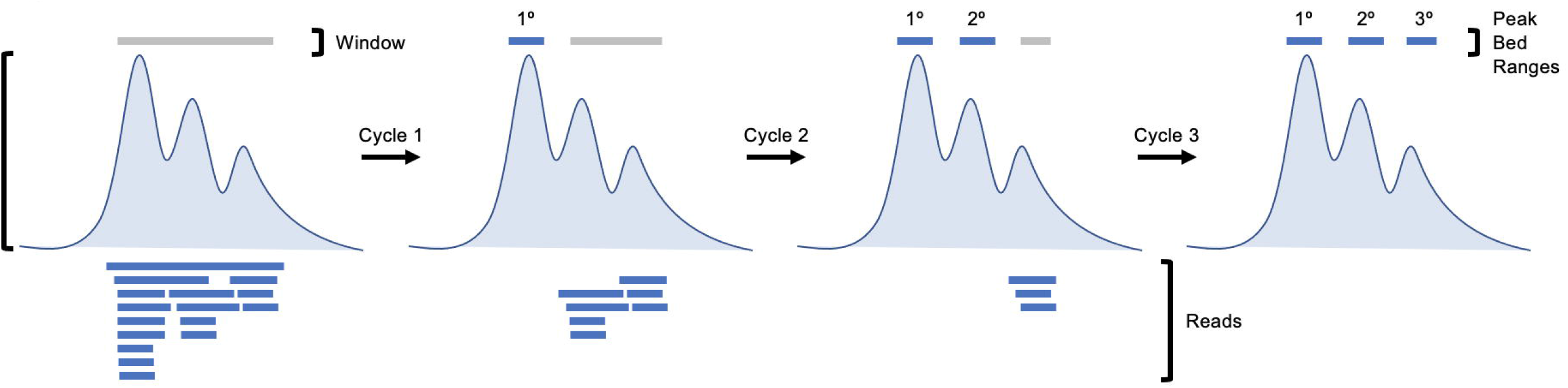
Diagram of repetitive peak calling by PACeR. Regions of the genome with 3 or more reads of coverage (windows) are identified and the region of maximal coverage within the window is determined (primary peak). Reads that fall under the primary peak are removed and new windows are identified and the region of maximal coverage within the new window is determined (secondary peak). Finally, reads that fall under the primary and secondary peaks are removed and new windows are identified and the region of maximal coverage within the new window is determined (tertiary peak). Peaks are then filtered to ensure that they are supported by reads from at least 2 libraries.

#### Motif Enrichment

Following peak identification, motif enrichment analysis is performed to determine which motifs are most abundant within peaks for a given sncRNA or sncRNA family. The center of the peak is determined and then expanded by an equal number of bases in either direction using BEDTools. Here we expand by 10 bases in both directions but this parameter can be adjusted by the user. The sequence of nucleotides within this region is then obtained using BEDTools and the number of peaks containing all possible 6- or 7-mer motifs is calculated.

### PACeR applied to chimeric RNA sequencing data from distinct preparatory approaches

To develop and validate PACeR, we tested the pipeline on datasets produced by two distinct molecular workflows, both of which sequenced Ago2-associated chimeric RNAs. We present the results of PACeR, first for analysis of a representative chimeric RNA sequencing dataset generated by our lab as described (MATERIALS AND METHODS) (Table 3, 5) and second for analysis of a benchmark published chimeric RNA sequencing dataset generated by the CLEAR-CLIP approach (Table 4, 5) (Moore et al. 2015). Uniquely mapped sncRNA-first chimeras were found to represent roughly 1.5 – 2.8% of post-FLASh reads, which is comparable to rates observed in similar analyses (Table 3) (Moore et al. 2015). As expected, libraries generated downstream of an immunoprecipitation (IP) step using an anti-Ago2 antibody (Sample1-4) produced significantly more chimeras than those prepared using isotype control IgG for the IP step (Control1-2) (Figure 4) and had a greater proportion of reads representing chimeras (Table 3). This supports that the chimeras detected with this method are Ago2-specific and reflect the interaction between sncRNAs and their targets within the RISC. For both our dataset and the previously published CLEAR-CLIP dataset we observed that the most enriched 6-mer motif within peaks called using this pipeline was the seed-matched binding site of the miRNA or miRNA family responsible for those peaks (Table 5).

**Table 3.** Summary of reads passing successive steps of PACeR pipeline for representative data.

| Sample | IP Antibody | R1/R2 Raw Reads | FLASH Reads | Cutadapt Reads | Deduplicated Reads | BLAST Aligned Reads | sncRNA-first Chimeras | Unique Aligned Reads | % of FLASH Reads |
| --- | --- | --- | --- | --- | --- | --- | --- | --- | --- |
| Sample1 | Ago2 | 93111817 | 82770147 | 40413104 | 29206799 | 6257678 | 3412974 | 1300235 | 1.57% |
| Sample2 | Ago2 | 138505593 | 128695154 | 79088139 | 53420319 | 11008923 | 6430642 | 2527163 | 1.96% |
| Sample3 | Ago2 | 61829813 | 47142440 | 25352183 | 19579823 | 4695142 | 2467235 | 1014930 | 2.15% |
| Sample4 | Ago2 | 69941142 | 61196930 | 45103518 | 33594273 | 7158176 | 4379156 | 1723694 | 2.82% |
| Control1 | IgG | 53979423 | 40705587 | 14932367 | 8088709 | 389236 | 173838 | 78185 | 0.19% |
| Control2 | IgG | 61851271 | 48311729 | 13298619 | 8013197 | 276492 | 118397 | 48549 | 0.10% |

**Table 4.** Summary of reads passing successive steps of PACeR pipeline for previously published data (Moore et al. 2015).

| Sample SRX ID | IP Antibody | Raw Reads | FLASH Reads | Cutadapt Reads | Deduplicated Reads | BLAST Aligned Reads | sncRNA-first Chimeras | Unique Aligned Reads | % of Raw Reads |
| --- | --- | --- | --- | --- | --- | --- | --- | --- | --- |
| SRX1247146 | pan-Ago | 8995221 | NA | 8251041 | 2196623 | 128589 | 59068 | 43410 | 0.48% |
| SRX1247147 | pan-Ago | 10019467 | NA | 8903130 | 1168755 | 83206 | 26025 | 8804 | 0.09% |
| SRX1247148 | pan-Ago | 10263108 | NA | 9445224 | 1899188 | 113572 | 47259 | 32675 | 0.32% |
| SRX1247151 | pan-Ago | 3804497 | NA | 3506694 | 1863882 | 213904 | 116560 | 47027 | 1.24% |
| SRX1247149* | pan-Ago | 9816796 | NA | 9064253 | 1905556 | 64234 | 4033 | 2421 | 0.02% |
| SRX1247150* | pan-Ago | 6615281 | NA | 6180788 | 1970629 | 64527 | 3720 | 2217 | 0.03% |

\*Controls prepared without the addition of ligase, see MATERIALS AND METHODS

**Table 5.**
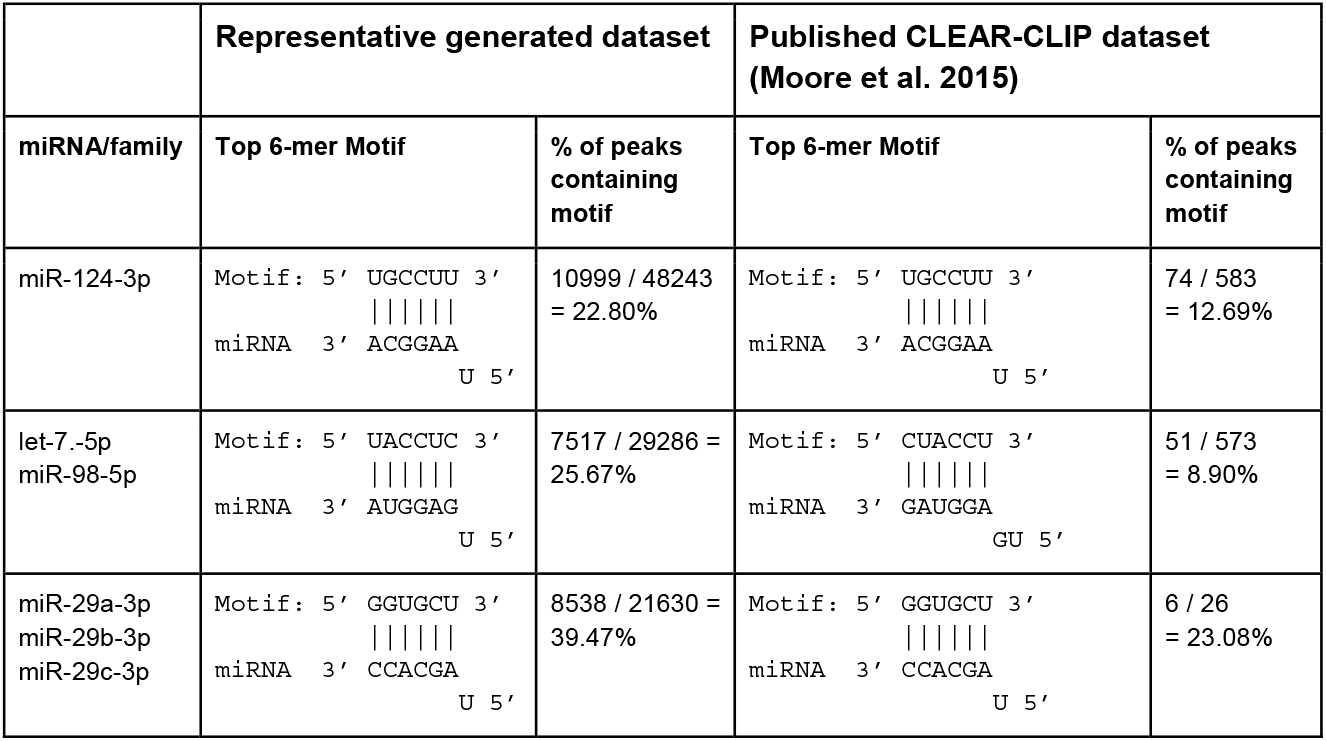
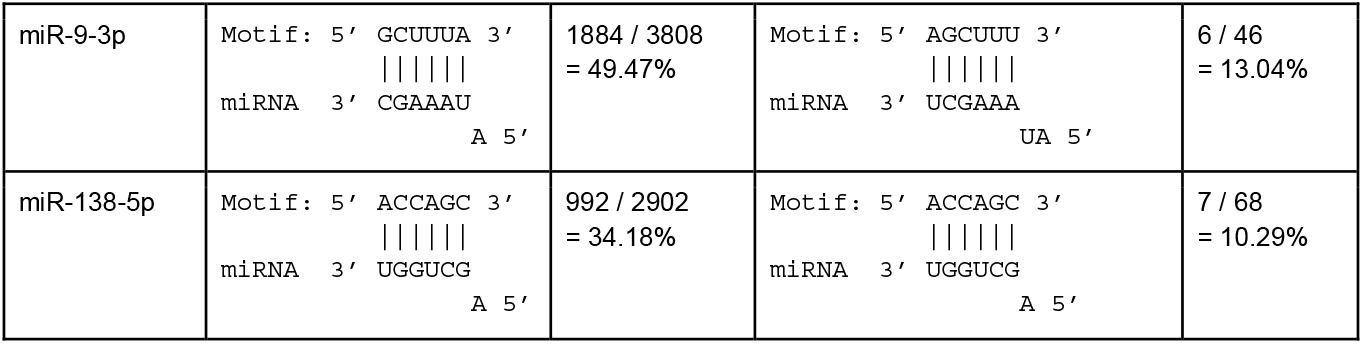
Motif enrichment within peaks identified for individual miRNAs of miRNA families.

**FIGURE 4.**
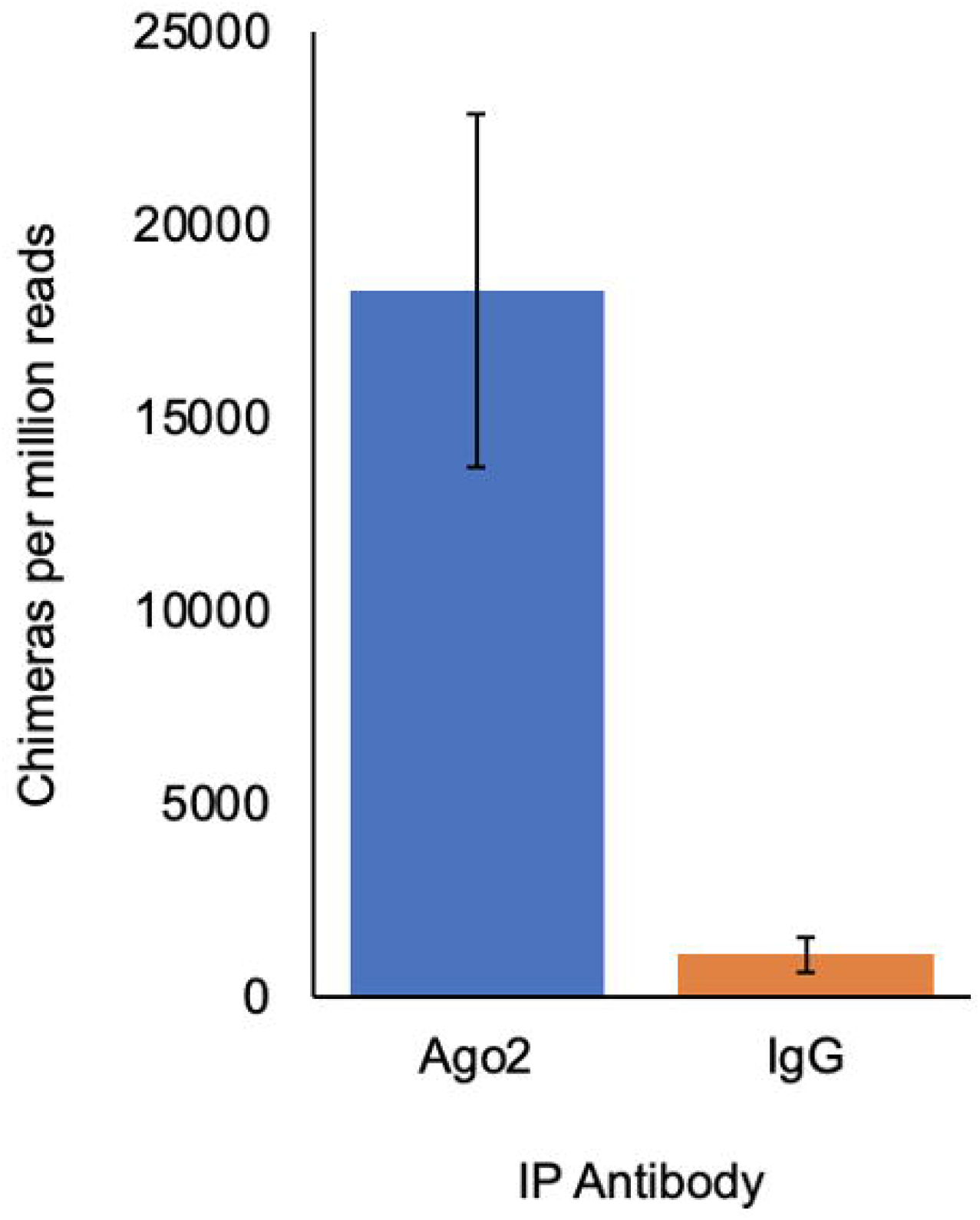
Chimeric sncRNA:target RNA reads generated using anti-Ago2 or IgG control antibody. Chimeric RNA sequencing libraries were generated with either anti-Ago2 (n=4) or isotype control IgG (n=2) antibodies for IP followed by library preparation and sequencing (150-base paired-end sequencing, 50-150 million reads per end). The number of chimeric RNA reads generated per million sequencing reads was calculated for IP with either Ago2 or isotype control IgG.

Using the representative dataset generated in our lab, peak calling using the PACeR pipeline can detect multiple interactions of the same sncRNA (or sncRNA family) within close proximity to one another, binding sites within introns, and binding sites within noncoding RNAs. The data presented below represent interactions between the let-7 family of miRNAs and various targets to demonstrate the ability of PACeR to accurately define miRNA target sites, especially when compared to other traditional peak callers.

### Adjacent miRNA binding sites

While miRNA binding sites are spread throughout the genome, multiple binding sites for a single miRNA or miRNA family are sometimes found in close proximity to one another which can increase their efficacy in downregulating targets (Saetrom et al. 2007). In initial attempts to identify miRNA binding sites from chimeric RNA sequencing data, we were unable to identify secondary miRNA binding sites that were in close proximity to primary sites. To rectify the issue, PACeR uses multiple cycles of peak calling, performed using reads that fall outside of the peak identified in the previous cycle(s). When considering the let-7-family of miRNAs in our representative data set, most additional sites were detected by 3 cycles of peak calling, as evidenced by the plateau in number of peaks identified (Figure 5).

**FIGURE 5.**
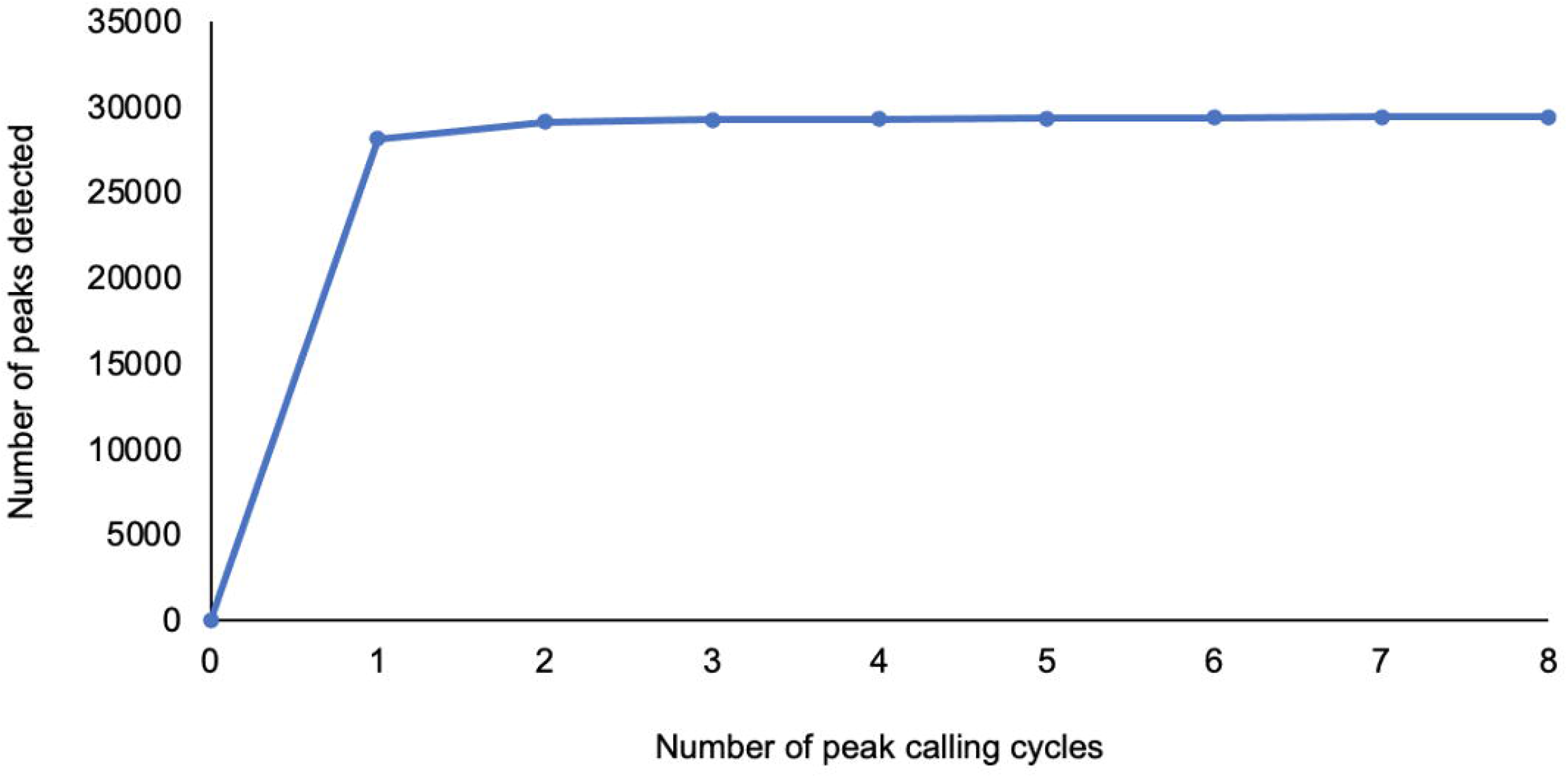
Number of peaks identified from chimeric RNA sequencing data following multiple rounds of peak calling for the let-7 family of miRNAs. Following a round of peak calling, reads that fell beneath the primary peak were removed and the remaining reads were again subjected to another round of peak calling. After all rounds of peak calling were complete, the peaks were filtered to require that they were supported by reads from at least 2 libraries.

As adjacent binding sites are not typically seen in CHIP- or CLIP-seq experiments, traditional peak callers are often unable to resolve proximal binding sites as two distinct sites. Figure 6 highlights one such example in which several traditional peak callers are unable to detect two proximal let-7 family binding sites within an exon of Speg in our representative data.

**FIGURE 6.**
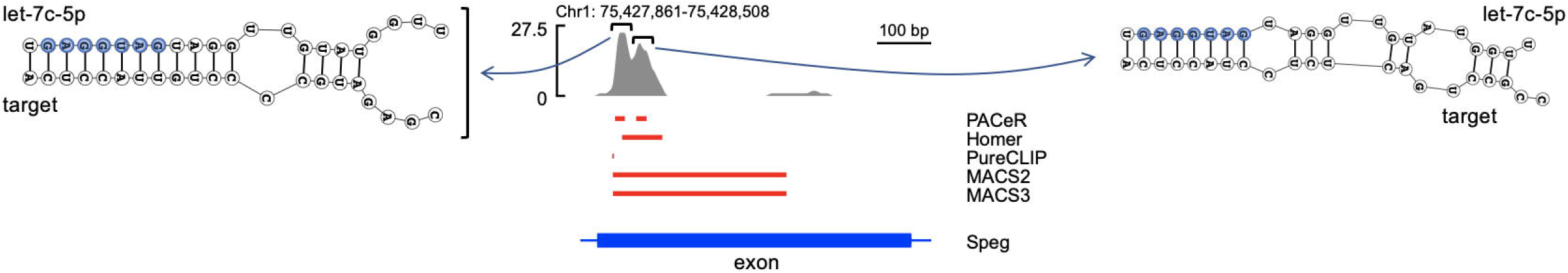
Adjacent miRNA binding sites resolved using PACeR. Two let-7 miRNA family proximal binding sites in an exon of Speg are often resolved as one site by traditional peak callers. The position of the peak identified by traditional peak callers is also skewed either before the peak (PureCLIP), between the two proximal sites (Homer), or in an area much larger than the peaks themselves (MACS2, MACS3). A representative binding interaction at this site is shown with let-7c-5p. Red bars represent genomic regions of peaks identified by each indicated peak caller. PACeR is able to identify the two distinct sites and restrict their positions to the respective peak maxima. Blue bar represents indicated gene regions. Genomic tracks were visualized with SparK (Kurtenbach and Harbour 2019). Binding interaction visualized using DuplexFold Web Server from the RNAstructure package (https://rna.urmc.rochester.edu/RNAstructureWeb/Servers/DuplexFold/DuplexFold.html, (Reuter and Mathews 2010)).

### MiRNA binding sites within introns

While the traditional view of miRNA binding to targets is that these binding sites occur within the 3’ UTR of target transcripts, there is growing evidence that miRNA-binding interactions can also occur in other regions of transcripts such as introns present in primary pre-mRNA (Sarshad et al. 2018). Figure 7 represents an example of a let-7 family binding site detected in our generated chimeric RNA-seq data within an annotated intron of Lrrtm4, a gene which encodes a transsynaptic adhesion protein (Chen et al. 2019; Reichman et al. 2020). It should be noted that such binding sites could actually reside in exons or other transcript regions due to alternatively spliced transcripts; accompanying total mRNA-seq data could be evaluated for comparison.

**FIGURE 7.**
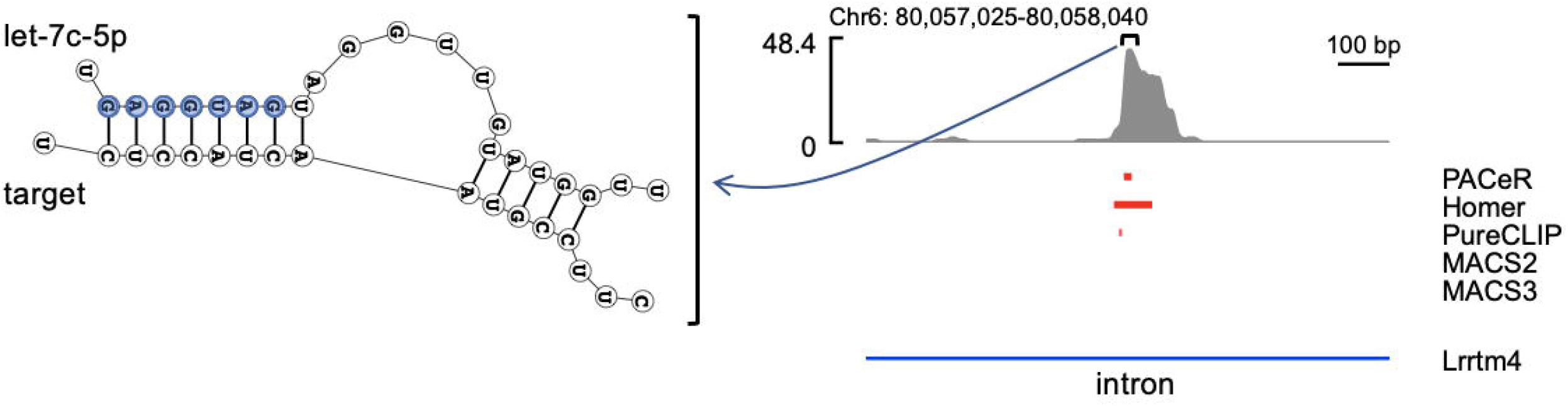
Identification of miRNA binding sites within introns. Red bars represent genomic regions of let-7 miRNA family binding peaks identified by each indicated peak caller; a peak is not identified by MACS2 or MACS3. A representative binding interaction at this site is shown with let-7c-5p. Blue bar represents indicated gene regions. Genomic tracks were visualized with SparK (Kurtenbach and Harbour, 2019). Binding interaction visualized using DuplexFold Web Server from the RNAstructure package (https://rna.urmc.rochester.edu/RNAstructureWeb/Servers/DuplexFold/DuplexFold.html (Reuter and Mathews 2010)).

### MiRNA binding site within a non-coding RNA

In addition to the canonical view of miRNA interactions with mRNAs, there is evidence that miRNAs can also interact with non-coding RNAs such as long non-coding RNAs (lncRNAs) (Yoon et al. 2014). Figure 8 illustrates an example of let-7 family miRNA binding sites detected in our generated chimeric RNA-seq data within MALAT1, a lncRNA that is highly expressed in the brain (Zhang et al. 2017) and whose stability has been shown to be regulated by miRNA binding (Leucci et al. 2013).

**FIGURE 8.**
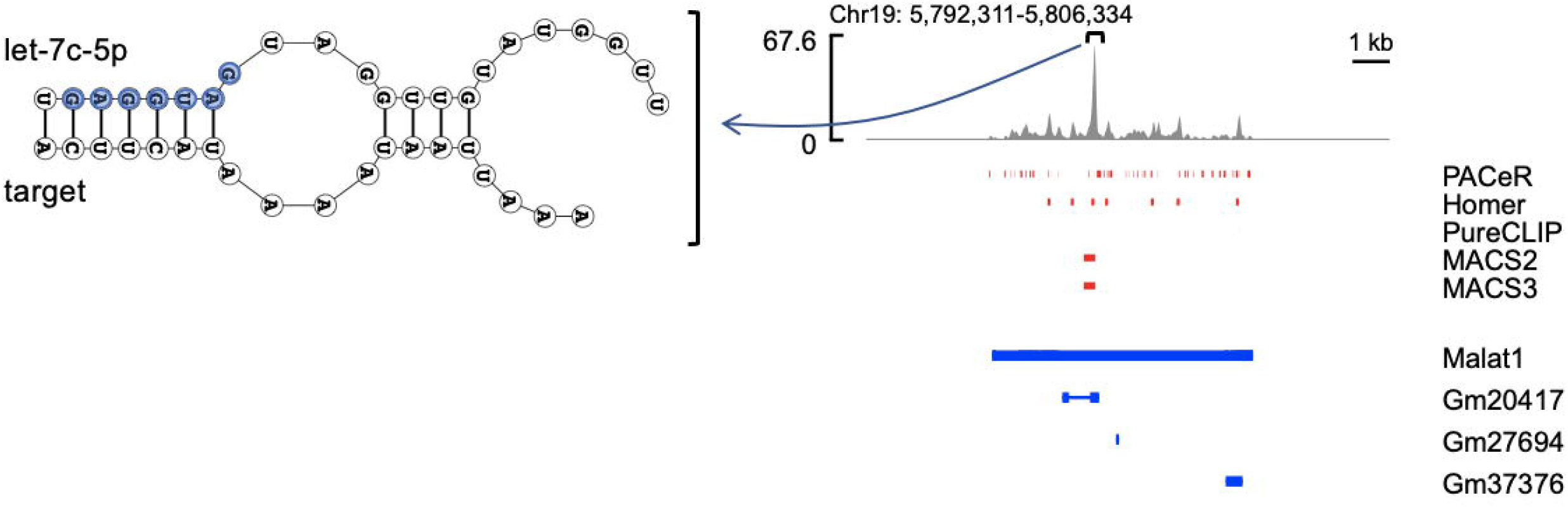
Identification of miRNA binding sites within non-coding RNAs. Red bars represent genomic regions of peaks identified by each indicated peak caller; PureCLIP is unable to identify the predominant peak. A representative binding interaction at this site is shown with let-7-5p. Blue bar represents indicated gene regions. Genomic tracks were visualized with SparK (Kurtenbach and Harbour, 2019). Binding interaction visualized using DuplexFold Web Server from the RNAstructure package (https://rna.urmc.rochester.edu/RNAstructureWeb/Servers/DuplexFold/DuplexFold.html, (Reuter and Mathews 2010)).

## Conclusions

### Considerations in the use of PACeR

PACeR uses several external packages/resources, such as Cutadapt, BLAST, and HISAT2. While these external packages were chosen with particular consideration given to their current broad public usage and maintenance, there may arise a situation when one or another of these external packages would become unreliable or for which there may be a better alternative for a user’s specific needs. The simplicity of the bash script and extensive annotation throughout should allow users to readily use other external packages as they require. However, to facilitate integration of packages other than those used in this script, users must ensure proper configuration of the input/output file formats for each step along the pipeline (e.g. FASTQ vs. FASTA files, BAM vs. BED files). The annotation included in the script will help users ensure the correct configuration, should an alternative external package be desirable.

### Limitations of PACeR

While PACeR is able to accurately identify miRNA binding sites in various settings that are often poorly characterized by other peak calling programs, such as adjacent binding sites, binding sites within introns, and binding sites within noncoding RNAs, there remain some sites that are difficult to properly define. For example, because miRNAs canonically interact with mature mRNAs, their binding sites may occur across exon-exon junctions. In its current configuration, the PACeR pipeline is unable to resolve these interactions as single binding sites and instead defines sites on both exons. While MACS3 is better able to define these interactions as single binding sites, this benefit seems outweighed by the major limitations described above. Additionally, the binding of miRNAs to exon-exon junctions is believed to be a rather infrequent event and would not significantly impact the results obtained from PACeR. However, users should consider this scenario when drawing conclusions from their data, particularly when considering miRNA binding sites within coding regions.

### Training

PACeR is a flexible tool that will allow users with minimal prior bioinformatics experience to readily customize analysis of their chimeric RNA libraries. This same flexibility might also have the propensity to lead to significant variation in the quality of analysis. Oversight and training in the appropriate selection of flexible variables, as well as record-keeping regarding these selections, will be a necessary step to ensure reproducible data.

The rapidly expanding field of miRNA biology has been greatly enhanced by the development of methods allowing for the unambiguous characterization of miRNA-target interactions. These methods also provide an avenue by which interactions between other sncRNAs and target RNAs may be described. However, a key bottleneck along the path to deriving meaningful conclusions from these methods has been the ability to process the chimeric RNA sequencing data. The chimeric RNA analysis pipeline presented here provides a ready-to-use, easy-to-understand way for scientists to escape this bottleneck and effectively process these data. This will significantly improve the field’s ability to understand the role that miRNAs and other sncRNAs play in post-transcriptional gene regulation and allow researchers to study how perturbations in these pathways impact normal cell biology and lead to disease.

### Ancillary Data Processing

After completing analysis of chimeric RNA-seq data with PACeR, users may wish to proceed with additional investigations beyond peak identification, such as comparisons of read coverage between biological conditions or determining the genes and pathways being targeted in their samples. BEDTools may be used to determine read coverage across peaks for individual samples after which differences between conditions may be readily assessed by employing traditional tools developed for differential “expression” analysis, such as DESeq2 (Love et al. 2014), to determine which peaks may be differentially bound between different experimental conditions. The output BED files from PACeR may be used with annotation packages such as Homer (http://homer.ucsd.edu/homer/index.html (Heinz et al. 2010)) to identify the genes and gene regions in which peaks reside. Additionally, gene ontology enrichment analysis may be performed on identified genes with tools such as The Gene Ontology Resource (http://geneontology.org/http://geneontology.org/ (Ashburner et al. 2000; The Gene Ontology Consortium 2021; Mi et al. 2019)). While not exhaustive, these examples illustrate possible ancillary data analyses that users may wish to perform downstream of outputs from the PACeR pipeline.

## MATERIALS AND METHODS

### Molecular approach for generating chimeric sncRNA:target RNA libraries

We provide an open-source pipeline which can be used for analyzing sequencing libraries of chimeric RNAs consisting of noncoding RNAs and target RNA pairs that are generated by multiple approaches (Table 1), with the intent of facilitating broader adoption of chimeric sequencing methods for studying the role of miRNAs and other sncRNAs in posttranscriptional gene regulation. Representative data to demonstrate the utility of this pipeline was generated by preparing chimeric RNA libraries using the following general approach (similar to (Bjerke and Yi 2020)). In brief, adult mouse forebrain tissue was homogenized and exposed to UV light (254 nm) to covalently crosslink RNAs to Ago. IP was carried out using an antibody against Ago2 (anti-Ago2 clone 2D4, Wako Pure Chemical Company, Catalog #018-22021) or IgG isotype control (Santa Cruz, Catalog #sc-2025) and bound RNAs were then trimmed with RNase before ligating the bound RNAs together (Figure 9A). After the formation of these chimeric RNAs, a DNA adapter was ligated to their 3’ end and the complexes were separated by SDS-PAGE and transferred to nitrocellulose. The bound RNAs were then released by Proteinase K treatment and an RNA adapter was ligated to their 5’ end. The chimeric RNAs with 5’ and 3’ adapters were reverse transcribed and amplified by PCR before sequencing. One important feature of the 5’ and 3’ adapters is the presence of random nucleotides at the ends that become ligated to the chimeric RNAs. These random nucleotides are used in the analysis pipeline to allow for multiple authentic interactions between the same sncRNA and the same target to be differentiated from PCR duplicates that may be generated during library preparation. The general structure of the generated chimeric RNAs is shown in Figure 9B. Libraries generated in our lab consisted of 4 chimeric sncRNA:target RNA libraries and 2 control libraries utilizing isotype control IgG in place of the Ago2 antibody during the IP step. These libraries were subjected to 150-base paired-end sequencing at a read depth between 50 and 150 million reads per end (Novogene, Beijing, China).

**FIGURE 9.**
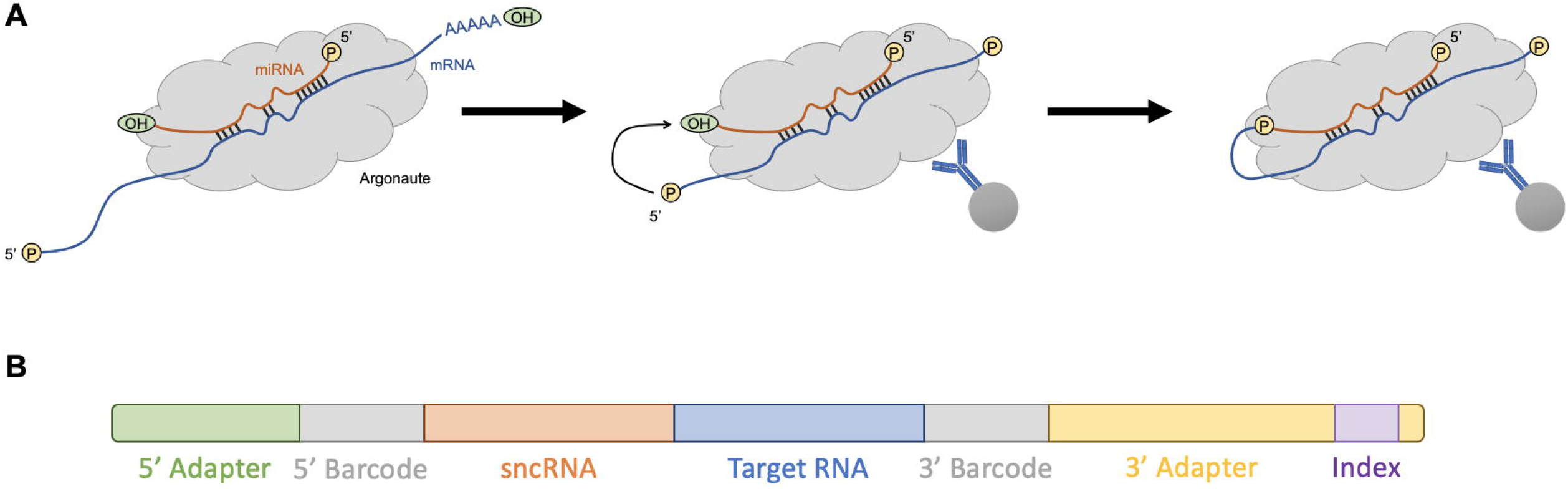
The generation of sncRNA:target chimeras and their features. (A) Following covalent crosslinking of RNAs to Argonaute via exposure to UV light, complexes are treated with an RNase to ensure that the 5’ end of the target RNA is in close proximity to the 3’ end of the miRNA. The two RNA strands within the Argonaute complex are then ligated together using a single-stranded RNA Ligase. ‘P’ denotes the terminal phosphate group on the RNA strand. (B) 5’ and 3’ adapters allow for annealing to the Illumina flow cell, 5’ and 3’ barcodes allow for the differentiation between multiple occurrences of the same sncRNA:target interaction and PCR duplicates, and the index sequence allows for multiple libraries to be pooled and run on the same flow cell.

### Pipeline use and implementation

Scripts for running the bioinformatic pipeline presented here (PACeR) are accessible on GitHub (https://github.com/Meffert-Lab/PACeR) and are summarized in Table 6. All analyses presented here were performed on a MacBook Pro (2017, 2.3 GHz Dual-Core Intel Core i5, 16 GB 2133 MHz LPDDR3, MacOS Monterey Version 12.2.1) using VirtualBox (Version 6.1.30) running Ubuntu (Version 20.04.3 LTS).

**Table 6.** Description of files available on GitHub (https://github.com/Meffert-Lab/PACeR).

| File Name | Description |
| --- | --- |
| 1_Pipeline_Setup_Tips.sh | Instructions for installing VirtualBox and Ubuntu, creating a shared folder between your computer and VirtualBox, installing Miniconda, and setting up a Miniconda environment with the requisite packages from running PACeR. |
| environment.yml | File for configuring the Miniconda environment with the requisite packages from running PACeR. |
| 2_Reference_File_Configuration.sh | Script for downloading and configuring the sncRNA reference list and the reference genome. |
| 3_Pipeline.sh | Script for processing compressed FASTQ files to generate a BAM file containing uniquely aligned reads annotated with the corresponding sncRNA within the read. |
| 3*_Pipeline_for_CLEAR-CLIP.sh | Script for processing compressed FASTQ files from a previously published dataset (Moore et al. 2015) to generate a BAM file containing uniquely aligned reads annotated with the corresponding sncRNA within the read. |
| 4a_Peak_Calling_Total.sh | Script for calling peaks from BAM files containing uniquely aligned reads annotated with the corresponding sncRNA within the read. |
| 4b_Peak_Calling_Subset.sh | Script for calling peaks from BAM files containing uniquely aligned reads annotated with the corresponding sncRNA within the read. Subsets reads by individual sncRNAs or sncRNA families and includes motif enrichment analysis. |

### Analysis of CLEAR-CLIP data

Chimeric RNA-seq data from a previously published benchmark dataset (Moore et al. 2015) were also processed by customizing variables in PACeR, (adapted version posted in Github as ‘3*_Pipeline_for_CLEAR-CLIP.sh’) to highlight the ability of the pipeline variables to be adjusted for chimeric RNA-seq datasets obtained using different biochemical methods. FASTQ files were obtained from the NCBI Sequence Read Archive (https://www.ncbi.nlm.nih.gov/sra) for biological replicates 1–6 (SRX1247146–SRX1247151). The reads were processed as described above with the following modifications according to details provided on the Gene Expression Omnibus (GEO) accession for the samples (example: https://www.ncbi.nlm.nih.gov/geo/query/acc.cgi?acc=GSM1881516): (1) no FLASh step performed as only single-end sequencing was performed; (2) during the Cutadapt step, the 3’ adapter sequence was changed to GTGTCAGTCACTTCCAGCGG, 5’ adapter sequence was changed to NNNNAGGGAGGACGATGCGG, and the requirement for linked adapters was removed; (3) the 5’ and 3’ barcode lengths were changed to 5 and 0, respectively. Peak calling and motif enrichment analysis was performed exactly as described above.

### Visualization of miRNA binding

The binding of miRNAs to their binding sites was visualized using the DuplexFold Web Server from the RNAstructure package (https://rna.urmc.rochester.edu/RNAstructureWeb/Servers/DuplexFold/DuplexFold.html, (Reuter and Mathews 2010)). The sequence of let-7c-5p from mouse (from miRBase (Kozomara et al. 2019); https://www.mirbase.org/) and the sequence underlying the peaks identified for the let-7 family were submitted to the DuplexFold Web Server and the predicted binding interactions were obtained.

## ACKNOWLEDGEMENTS

We thank members of the Meffert laboratory for helpful discussions, including B. Powell and S. Eadara for critical feedback on the pipeline. Funding: This work was supported by a Braude Foundation award and Blaustein Endowment for Pain Research and Education award to M.K.M., and National Institutes of Health (NIH) grant F31MH124282 to W.T.M.

## REFERENCES

Altschul SF, Gish W, Miller W, Myers EW, Lipman DJ. 1990. Basic local alignment search tool. J Mol Biol 215: 403–410.

Andrews S. 2010. FastQC: A Quality Control Tool for High Throughput Sequence Data. http://www.bioinformatics.babraham.ac.uk/projects/fastqc/ (Accessed March 17, 2022).

Ashburner M, Ball CA, Blake JA, Botstein D, Butler H, Cherry JM, Davis AP, Dolinski K, Dwight SS, Eppig JT, et al. 2000. Gene Ontology: tool for the unification of biology. Nat Genet 25: 25–29.

Auwera GAV der, O’Connor BD. 2020. Genomics in the Cloud: Using Docker, GATK, and WDL in Terra. 1st ed. O’Reilly Media, Sebastopol, CA.

Baruzzo G, Hayer KE, Kim EJ, Di Camillo B, FitzGerald GA, Grant GR. 2017. Simulation-based comprehensive benchmarking of RNA-seq aligners. Nat Methods 14: 135–139.

Bazzini AA, Lee MT, Giraldez AJ. 2012. Ribosome profiling shows that miR-430 reduces translation before causing mRNA decay in zebrafish. Science 336: 233–237.

Béthune J, Artus-Revel CG, Filipowicz W. 2012. Kinetic analysis reveals successive steps leading to miRNA-mediated silencing in mammalian cells. EMBO Rep 13: 716–723.

Bjerke GA, Yi R. 2020. Integrated analysis of directly captured microRNA targets reveals the impact of microRNAs on mammalian transcriptome. RNA 26: 306–323.

Broughton JP, Lovci MT, Huang JL, Yeo GW, Pasquinelli AE. 2016. Pairing beyond the Seed Supports MicroRNA Targeting Specificity. Mol Cell 64: 320–333.

Broughton JP, Pasquinelli AE. 2013. Identifying Argonaute binding sites in Caenorhabditis elegans using iCLIP. Methods 63: 119–125.

Chen Y-C, Liu Y-L, Tsai S-J, Kuo P-H, Huang S-S, Lee Y-S. 2019. LRRTM4 and PCSK5 Genetic Polymorphisms as Markers for Cognitive Impairment in A Hypotensive Aging Population: A Genome-Wide Association Study in Taiwan. J Clin Med 8.

Chipman LB, Pasquinelli AE. 2019. miRNA Targeting: Growing beyond the Seed. Trends Genet 35: 215–222.

Chi SW, Zang JB, Mele A, Darnell RB. 2009. Argonaute HITS-CLIP decodes microRNA-mRNA interaction maps. Nature 460: 479–486.

Ciafrè SA, Galardi S. 2013. microRNAs and RNA-binding proteins: a complex network of interactions and reciprocal regulations in cancer. RNA Biol 10: 935–942.

Djuranovic S, Nahvi A, Green R. 2012. miRNA-mediated gene silencing by translational repression followed by mRNA deadenylation and decay. Science 336: 237–240.

Duan Y, Veksler-Lublinsky I, Ambros V. 2022. Critical contribution of 3’ non-seed base pairing to the in vivo function of the evolutionarily conserved let-7a microRNA. Cell Rep 39: 110745.

Dweep H, Sticht C, Pandey P, Gretz N. 2011. miRWalk--database: prediction of possible miRNA binding sites by “walking” the genes of three genomes. J Biomed Inform 44: 839–847.

Enright AJ, John B, Gaul U, Tuschl T, Sander C, Marks DS. 2003. MicroRNA targets in Drosophila. Genome Biol 5: R1.

Ewels P, Magnusson M, Lundin S, Käller M. 2016. MultiQC: summarize analysis results for multiple tools and samples in a single report. Bioinformatics 32: 3047–3048.

Fabian MR, Mathonnet G, Sundermeier T, Mathys H, Zipprich JT, Svitkin YV, Rivas F, Jinek M, Wohlschlegel J, Doudna JA, et al. 2009. Mammalian miRNA RISC recruits CAF1 and PABP to affect PABP-dependent deadenylation. Mol Cell 35: 868–880.

Forman JJ, Legesse-Miller A, Coller HA. 2008. A search for conserved sequences in coding regions reveals that the let-7 microRNA targets Dicer within its coding sequence. Proc Natl Acad Sci USA 105: 14879–14884.

Friedman RC, Farh KK-H, Burge CB, Bartel DP. 2009. Most mammalian mRNAs are conserved targets of microRNAs. Genome Res 19: 92–105.

Gay LA, Sethuraman S, Thomas M, Turner PC, Renne R. 2018. Modified Cross-Linking, Ligation, and Sequencing of Hybrids (qCLASH) Identifies Kaposi’s Sarcoma-Associated Herpesvirus MicroRNA Targets in Endothelial Cells. J Virol 92.

Gregory RI, Yan K-P, Amuthan G, Chendrimada T, Doratotaj B, Cooch N, Shiekhattar R. 2004. The Microprocessor complex mediates the genesis of microRNAs. Nature 432: 235–240.

Grosswendt S, Filipchyk A, Manzano M, Klironomos F, Schilling M, Herzog M, Gottwein E, Rajewsky N. 2014. Unambiguous identification of miRNA:target site interactions by different types of ligation reactions. Mol Cell 54: 1042–1054.

Guan L, Karaiskos S, Grigoriev A. 2020. Inferring targeting modes of Argonaute-loaded tRNA fragments. RNA Biol 17: 1070–1080.

Hafner M, Landthaler M, Burger L, Khorshid M, Hausser J, Berninger P, Rothballer A, Ascano M, Jungkamp A-C, Munschauer M, et al. 2010. Transcriptome-wide identification of RNA-binding protein and microRNA target sites by PAR-CLIP. Cell 141: 129–141.

Hafner M, Lianoglou S, Tuschl T, Betel D. 2012. Genome-wide identification of miRNA targets by PAR-CLIP. Methods 58: 94–105.

Hammell CM, Lubin I, Boag PR, Blackwell TK, Ambros V. 2009. nhl-2 Modulates microRNA activity in Caenorhabditis elegans. Cell 136: 926–938.

Han J, Lee Y, Yeom K-H, Kim Y-K, Jin H, Kim VN. 2004. The Drosha-DGCR8 complex in primary microRNA processing. Genes Dev 18: 3016–3027.

Heinz S, Benner C, Spann N, Bertolino E, Lin YC, Laslo P, Cheng JX, Murre C, Singh H, Glass CK. 2010. Simple combinations of lineage-determining transcription factors prime cis-regulatory elements required for macrophage and B cell identities. Mol Cell 38: 576–589.

Helwak A, Kudla G, Dudnakova T, Tollervey D. 2013. Mapping the human miRNA interactome by CLASH reveals frequent noncanonical binding. Cell 153: 654–665.

Hu W, Coller J. 2012. What comes first: translational repression or mRNA degradation? The deepening mystery of microRNA function. Cell Res 22: 1322–1324.

Johnston M, Hutvagner G. 2011. Posttranslational modification of Argonautes and their role in small RNA-mediated gene regulation. Silence 2: 5.

John B, Enright AJ, Aravin A, Tuschl T, Sander C, Marks DS. 2004. Human MicroRNA targets. PLoS Biol 2: e363.

Ketting RF, Fischer SE, Bernstein E, Sijen T, Hannon GJ, Plasterk RH. 2001. Dicer functions in RNA interference and in synthesis of small RNA involved in developmental timing in C. elegans. Genes Dev 15: 2654–2659.

Kim D, Paggi JM, Park C, Bennett C, Salzberg SL. 2019. Graph-based genome alignment and genotyping with HISAT2 and HISAT-genotype. Nat Biotechnol 37: 907–915.

Kiriakidou M, Nelson PT, Kouranov A, Fitziev P, Bouyioukos C, Mourelatos Z, Hatzigeorgiou A. 2004. A combined computational-experimental approach predicts human microRNA targets. Genes Dev 18: 1165–1178.

Knight SW, Bass BL. 2001. A role for the RNase III enzyme DCR-1 in RNA interference and germ line development in Caenorhabditis elegans. Science 293: 2269–2271.

Kozomara A, Birgaoanu M, Griffiths-Jones S. 2019. miRBase: from microRNA sequences to function. Nucleic Acids Res 47: D155–D162.

Krakau S, Richard H, Marsico A. 2017. PureCLIP: capturing target-specific protein-RNA interaction footprints from single-nucleotide CLIP-seq data. Genome Biol 18: 240.

Krek A, Grün D, Poy MN, Wolf R, Rosenberg L, Epstein EJ, MacMenamin P, da Piedade I, Gunsalus KC, Stoffel M, et al. 2005. Combinatorial microRNA target predictions. Nat Genet 37: 495–500.

Kumar P, Anaya J, Mudunuri SB, Dutta A. 2014. Meta-analysis of tRNA derived RNA fragments reveals that they are evolutionarily conserved and associate with AGO proteins to recognize specific RNA targets. BMC Biol 12: 78.

Kumar P, Mudunuri SB, Anaya J, Dutta A. 2015. tRFdb: a database for transfer RNA fragments. Nucleic Acids Res 43: D141–5.

Kurtenbach S, Harbour JW. 2019. SparK: A Publication-quality NGS Visualization Tool. BioRxiv.

Kuscu C, Kumar P, Kiran M, Su Z, Malik A, Dutta A. 2018. tRNA fragments (tRFs) guide Ago to regulate gene expression post-transcriptionally in a Dicer-independent manner. RNA 24: 1093–1105.

Lal A, Navarro F, Maher CA, Maliszewski LE, Yan N, O’Day E, Chowdhury D, Dykxhoorn DM, Tsai P, Hofmann O, et al. 2009. miR-24 Inhibits cell proliferation by targeting E2F2, MYC, and other cell-cycle genes via binding to “seedless” 3’UTR microRNA recognition elements. Mol Cell 35: 610–625.

Lee RC, Feinbaum RL, Ambros V. 1993. The C. elegans heterochronic gene lin-4 encodes small RNAs with antisense complementarity to lin-14. Cell 75: 843–854.

Lee Y, Ahn C, Han J, Choi H, Kim J, Yim J, Lee J, Provost P, Rådmark O, Kim S, et al. 2003. The nuclear RNase III Drosha initiates microRNA processing. Nature 425: 415–419.

Lee Y, Jeon K, Lee J-T, Kim S, Kim VN. 2002. MicroRNA maturation: stepwise processing and subcellular localization. EMBO J 21: 4663–4670.

Lee Y, Kim M, Han J, Yeom K-H, Lee S, Baek SH, Kim VN. 2004. MicroRNA genes are transcribed by RNA polymerase II. EMBO J 23: 4051–4060.

Leucci E, Patella F, Waage J, Holmstrøm K, Lindow M, Porse B, Kauppinen S, Lund AH. 2013. microRNA-9 targets the long non-coding RNA MALAT1 for degradation in the nucleus. Sci Rep 3: 2535.

Lewis BP, Burge CB, Bartel DP. 2005. Conserved seed pairing, often flanked by adenosines, indicates that thousands of human genes are microRNA targets. Cell 120: 15–20.

Lewis BP, Shih I, Jones-Rhoades MW, Bartel DP, Burge CB. 2003. Prediction of mammalian microRNA targets. Cell 115: 787–798.

Long D, Lee R, Williams P, Chan CY, Ambros V, Ding Y. 2007. Potent effect of target structure on microRNA function. Nat Struct Mol Biol 14: 287–294.

Love MI, Huber W, Anders S. 2014. Moderated estimation of fold change and dispersion for RNA-seq data with DESeq2. Genome Biol 15: 550.

Lytle JR, Yario TA, Steitz JA. 2007. Target mRNAs are repressed as efficiently by microRNA-binding sites in the 5’ UTR as in the 3’ UTR. Proc Natl Acad Sci USA 104: 9667–9672.

Magoč T, Salzberg SL. 2011. FLASH: fast length adjustment of short reads to improve genome assemblies. Bioinformatics 27: 2957–2963.

Martin M. 2011. Cutadapt removes adapter sequences from high-throughput sequencing reads. EMBnet j 17: 10.

Matranga C, Tomari Y, Shin C, Bartel DP, Zamore PD. 2005. Passenger-strand cleavage facilitates assembly of siRNA into Ago2-containing RNAi enzyme complexes. Cell 123: 607–620.

Meijer HA, Kong YW, Lu WT, Wilczynska A, Spriggs RV, Robinson SW, Godfrey JD, Willis AE, Bushell M. 2013. Translational repression and eIF4A2 activity are critical for microRNA-mediated gene regulation. Science 340: 82–85.

Mishima Y, Fukao A, Kishimoto T, Sakamoto H, Fujiwara T, Inoue K. 2012. Translational inhibition by deadenylation-independent mechanisms is central to microRNA-mediated silencing in zebrafish. Proc Natl Acad Sci USA 109: 1104–1109.

Miyoshi K, Tsukumo H, Nagami T, Siomi H, Siomi MC. 2005. Slicer function of Drosophila Argonautes and its involvement in RISC formation. Genes Dev 19: 2837–2848.

Mi H, Muruganujan A, Ebert D, Huang X, Thomas PD. 2019. PANTHER version 14: more genomes, a new PANTHER GO-slim and improvements in enrichment analysis tools. Nucleic Acids Res 47: D419–D426.

Moore MJ, Scheel TKH, Luna JM, Park CY, Fak JJ, Nishiuchi E, Rice CM, Darnell RB. 2015. miRNA-target chimeras reveal miRNA 3’-end pairing as a major determinant of Argonaute target specificity. Nat Commun 6: 8864.

Moore MJ, Zhang C, Gantman EC, Mele A, Darnell JC, Darnell RB. 2014. Mapping Argonaute and conventional RNA-binding protein interactions with RNA at single-nucleotide resolution using HITS-CLIP and CIMS analysis. Nat Protoc 9: 263–293.

Moretti F, Kaiser C, Zdanowicz-Specht A, Hentze MW. 2012. PABP and the poly(A) tail augment microRNA repression by facilitated miRISC binding. Nat Struct Mol Biol 19: 603–608.

Quinlan AR, Hall IM. 2010. BEDTools: a flexible suite of utilities for comparing genomic features. Bioinformatics 26: 841–842.

Reichman RD, Gaynor SC, Monson ET, Gaine ME, Parsons MG, Zandi PP, Potash JB, Willour VL. 2020. Targeted sequencing of the LRRTM gene family in suicide attempters with bipolar disorder. Am J Med Genet B Neuropsychiatr Genet 183: 128–139.

Reuter JS, Mathews DH. 2010. RNAstructure: software for RNA secondary structure prediction and analysis. BMC Bioinformatics 11: 129.

Ricci EP, Limousin T, Soto-Rifo R, Rubilar PS, Decimo D, Ohlmann T. 2013. miRNA repression of translation in vitro takes place during 43S ribosomal scanning. Nucleic Acids Res 41: 586–598.

Saetrom P, Heale BSE, Snøve O, Aagaard L, Alluin J, Rossi JJ. 2007. Distance constraints between microRNA target sites dictate efficacy and cooperativity. Nucleic Acids Res 35: 2333–2342.

Sarshad AA, Juan AH, Muler AIC, Anastasakis DG, Wang X, Genzor P, Feng X, Tsai P-F, Sun H-W, Haase AD, et al. 2018. Argonaute-miRNA Complexes Silence Target mRNAs in the Nucleus of Mammalian Stem Cells. Mol Cell 71: 1040-1050.e8.

Shah A, Qian Y, Weyn-Vanhentenryck SM, Zhang C. 2017. CLIP Tool Kit (CTK): a flexible and robust pipeline to analyze CLIP sequencing data. Bioinformatics 33: 566–567.

Stark A, Brennecke J, Russell RB, Cohen SM. 2003. Identification of Drosophila MicroRNA targets. PLoS Biol 1: E60.

Tay Y, Zhang J, Thomson AM, Lim B, Rigoutsos I. 2008. MicroRNAs to Nanog, Oct4 and Sox2 coding regions modulate embryonic stem cell differentiation. Nature 455: 1124–1128.

The Gene Ontology Consortium. 2021. The Gene Ontology resource: enriching a GOld mine. Nucleic Acids Res 49: D325–D334.

Wang B, Bao L. 2017. Axonal microRNAs: localization, function and regulatory mechanism during axon development. J Mol Cell Biol 9: 82–90.

Wang J-H, Chen W-X, Mei S-Q, Yang Y-D, Yang J-H, Qu L-H, Zheng L-L. 2022. tsRFun: a comprehensive platform for decoding human tsRNA expression, functions and prognostic value by high-throughput small RNA-Seq and CLIP-Seq data. Nucleic Acids Res 50: D421–D431.

Wightman B, Ha I, Ruvkun G. 1993. Posttranscriptional regulation of the heterochronic gene lin-14 by lin-4 mediates temporal pattern formation in C. elegans. Cell 75: 855–862.

Xiao Q, Gao P, Huang X, Chen X, Chen Q, Lv X, Fu Y, Song Y, Wang Z. 2021. tRFTars: predicting the targets of tRNA-derived fragments. J Transl Med 19: 88.

Yoon J-H, Abdelmohsen K, Gorospe M. 2014. Functional interactions among microRNAs and long noncoding RNAs. Semin Cell Dev Biol 34: 9–14.

Zdanowicz A, Thermann R, Kowalska J, Jemielity J, Duncan K, Preiss T, Darzynkiewicz E, Hentze MW. 2009. Drosophila miR2 primarily targets the m7GpppN cap structure for translational repression. Mol Cell 35: 881–888.

Zhang C, Darnell RB. 2011. Mapping in vivo protein-RNA interactions at single-nucleotide resolution from HITS-CLIP data. Nat Biotechnol 29: 607–614.

Zhang X, Hamblin MH, Yin K-J. 2017. The long noncoding RNA Malat1: Its physiological and pathophysiological functions. RNA Biol 14: 1705–1714.

Zhang Y, Liu T, Meyer CA, Eeckhoute J, Johnson DS, Bernstein BE, Nusbaum C, Myers RM, Brown M, Li W, et al. 2008. Model-based analysis of ChIP-Seq (MACS). Genome Biol 9: R137.

Zuo Y, Zhu L, Guo Z, Liu W, Zhang J, Zeng Z, Wu Q, Cheng J, Fu X, Jin Y, et al. 2021. tsRBase: a comprehensive database for expression and function of tsRNAs in multiple species. Nucleic Acids Res 49: D1038–D1045.

